# TOWARDS QUANTITATIVE FORECASTING OF SPECIES YIELDS WITH INCOMPLETE INFORMATION ON MODEL PARAMETERS

**DOI:** 10.1101/2021.03.25.437089

**Authors:** Hugo Fort

## Abstract

Predicting both the absolute and the relative abundance of species in a spatial patch is of paramount interest in areas like, agriculture, ecology and environmental science.

The linear Lotka-Volterra generalized equations (LLVGE) serve for describing the dynamics of communities of species connected by negative as well as positive interspecific interactions.

Here we particularize these LLVGE to the case of single trophic ecological communities, like mixtures of plants, with *S* >2 species. Thus, by estimating the LLVGE parameters from the yields in monoculture and biculture experiments, the LLVGE are able to produce decently accurate predictions for species yields.

However, a common situation we face is that we don’t know all the parameters appearing in the LLVGE. Indeed, for large values of *S*, only a fraction of the experiments necessary for estimating the model parameters is commonly carried out. We then analyze which quantitative predictions are possible with an incomplete knowledge of the parameters.

## 1 INTRODUCTION

The ability to predict the absolute and the relative abundance of species in a place is of paramount interest in areas like, agriculture, ecology and environmental science. Indeed, abundance is an ecological quantity of crucial importance when making management and conservation decisions (Andrewartha and Birch 1954; Krebs 1978; Gaston 1994). For species of value to humans, estimates of their future yields will be essential for regulating current and future harvests. For species of conservation concern, their future populations will be an important determinant of their extinction risk.

Interspecific interactions strongly influence the absolute and relative abundances of species within a patch (Sousa 1984). In the case of agricultural crops, the mathematical methods commonly employed to forecast the yield have been mostly linear statistical models (NASS-SMB 2012). However, the interest in phenomenological modeling of biomass production and yield of crops has increased recently (Marcelis et al. 1998; Kirwan et al. 2009; Halty et al. 2017). Phenomenological models have in general a short computing time and they usually contain few state variables, making them helpful tools for reliable predictions.

Here we start by briefly reviewing the linear Lotka-Volterra *generalized* equations (LLVGE) (Pastor 2008; Fort 2020a) which serve for describing the dynamics of communities of species connected by negative as well as positive interspecific interactions. More specifically, we particularize these LLVGE to the case of a single trophic level community with *S* >2 species, either artificial or natural.

However, a common situation we face is that we don’t know all the parameters appearing in the LLVGE. Therefore, we analyze which quantitative predictions are possible with an incomplete knowledge of the parameters. We discuss two approximations that allow using these LLVGE as a quantitative tool.

## 2 THE LOTKA- VOLTERRA GENERALIZED LINEAR MODEL

If *n_i_* denotes the *yield* of species *i* (e.g. its density or its biomass density) the LLVCE for *S* coexisting species consist in a set of linear equations for the per capita growth rates 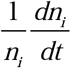 given by:

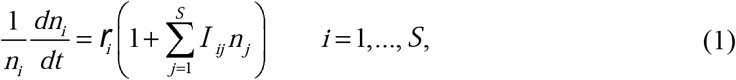

where 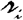 is the intrinsic growth rate of species *i*, with dimension of time^−1^, and *I_ij_* is an interaction coefficient, with dimension of yield^−1^, quantifying the *per capita* effect of species *j* on species *i* through their pairwise interaction. This is a phenomenological model since it describes how the abundance of one species affects the abundance of another, without specifically including a particular mechanism for such interaction (Morin 2011). ‘Generalized’ means that all the combinations of pairs of signs for both species (−/−,−/+,−/0, +/0, +/+) are possible.

### 2.1 The linear Lotka-Volterra generalized equations for single trophic communities

In the case of a single trophic community, the trophic level below the one of the community (e.g. plants for a community of herbivore species), corresponding to the food required for individuals of a species *i* to thrive, is modeled through a parameter 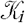 known as the *carrying capacity* (Pastor 2008). 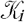 is interpreted as the maximum sustainable yield for species *i* in monoculture, *i.e*. when the species is isolated from the rest of the other species making up the community. It turns out that *I_ij_* can be thus written in terms of a non-dimensional interaction coefficient *α_ij_* (Fort 2020):

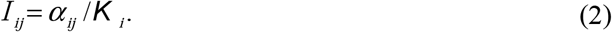

It is customary to take the intraspecific competition coefficients *α_ii_* = −1. Hence, by (1) and (2) the LLVGE become:

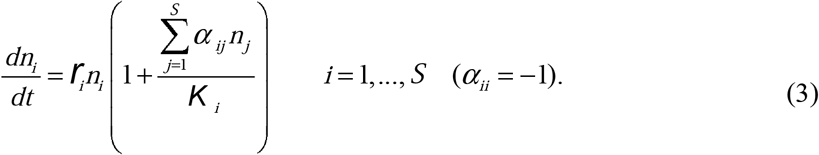

### 2.2 Obtaining the model parameters from monoculture and biculture experiments

A straightforward procedure to get the set of model parameters 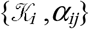 is to perform, until the equilibrium state is attained: a) the *S* single species or monoculture experiments, and from each of them to estimate the carrying capacities as the yield of the species *i* in monoculture 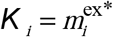 (the superscript ‘ex’ is to emphasize that this is an experimentally measured quantity and the star denotes equilibrium); b) the *S*×(*S*-1)/2 pairwise experiments producing the *biculture* yields, 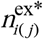 and 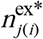 (the subscripts *i*(*j*) and *j*(*i*) stand for the yield of species *i* in presence of species *j* and *vice ver*sa). We then obtain *α_ij_* and *α_ji_*, as (Fort 2020a):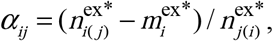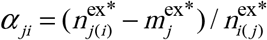.

## 3 WORKING WITH IMPERFECT INFORMATION

A main limitation of the above experimental procedure is that it is only feasible provided *S* is not too large. This is because the number of required experiments grows as *S*^2^. Thus, for large values of *S*, only a fraction of these experiments is commonly carried out and consequently we have an incomplete knowledge of the *S*^2^ parameters required to compute the equilibrium yields from the LLVGE.

However, there are situations in which we can still predict some relevant quantities about species yields with an incomplete knowledge of the LLVGE parameters. With this aim, it is convenient to work with relative yields

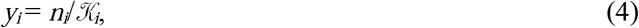

*i.e.* the species yield in mixture normalized by its yield in monoculture. It turns out that the equilibrium^1^ relative yields verify (Fort 2020a):

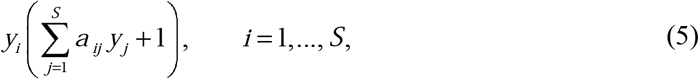

where

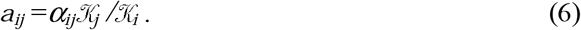

Notice thus that the diagonal elements of the matrix **A**=[*a_ij_*], corresponding to intra-specific competition, remain equal to −1 since 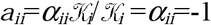.

Let us present a couple of recently introduced methods for making quantitative predictions with incomplete information on the matrix **A**, motivating each method through a different practical problem in crop science.

First, we can think about mixtures of crops grown for biomass, forage, or food production, and the related phenomenon of *overyielding*, i.e. the increased biomass production in species mixtures relative to monoculture (Beckage & Gross 2006). E.g., a popular aggregate metric used to quantify the overyielding of diverse crop mixtures relative to crop monocultures is the Relative Yield Total (*RYT*) (de Wit 1970):

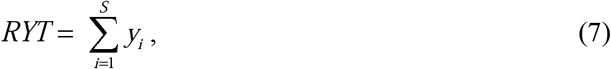

A *RYT* > 1 implies that the yield performance will be better in polyculture than in monoculture (Vandermeer 1989). Another index directly connected with the *RYT* is the Mean Relative Yield (*MRY*),

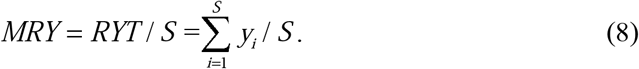

In this case we can use the recently proposed *mean field* approximation (MFA) (Fort 2018b;Fort & Segura 2018; Fort 2020a). The name alludes to the resemblance of this approximation with the one commonly used in physics, consisting in replacing spatially dependent variables by a constant equal to their mean value. Here the average will be taken over the values of the interspecific interaction coefficients. That is, the off-diagonal part of the interaction matrix [*a_ij_*] is replaced by a “mean field” competition coefficient given by the average over the sample of available interspecific competition coefficients:

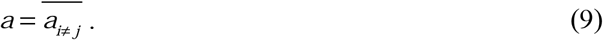

In such a way we get a *mean field matrix*:

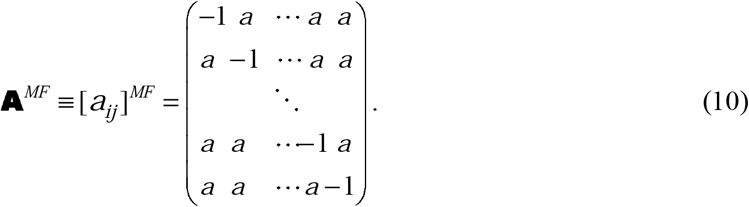

The MFA allows expressing the *RYT* and *MRY* as a simple functions of *a* and *S* (Fort 2018b, Fort 2020a):

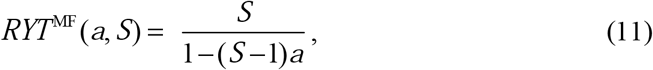

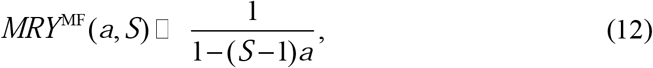

Thus the MFA, through Eqs. (11) and (12), is particularly suited to address an important question in ecology, namely, how the total or mean relative yield depends with the species richness and the mean intensity of competition. Comparisons between predictions generated by formula (11) against data from a dataset of 25 experiments involving *S* > 2 species and *a* < 0 are shown in Fig. 1 as a function of both *S* and *a* (Fort 2018b).

**Figure 1.**
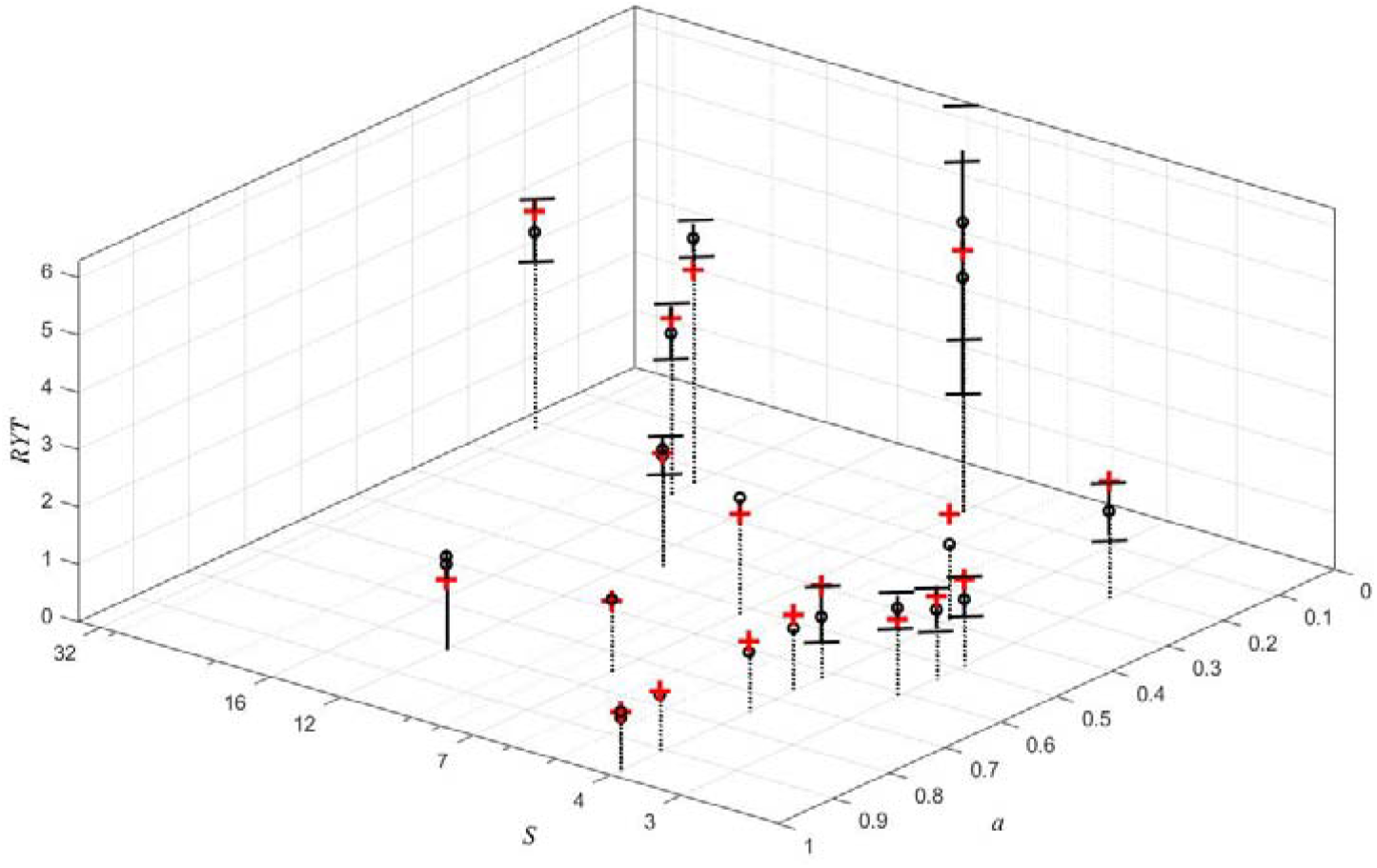
Empirical (o) and theoretical *RYT*, predicted by the MFA Eq. (11) (red crosses), for a set of 25 experiments (see text) as a function of *S* (log scale) and the mean competition parameter *a*. The experimental error bars correspond to ± the standard error (SE).

Second, suppose one is interested in predicting the yield of a *particular* species embedded in a community with other species belonging to the same trophic level. *E.g*. a crop coexisting with different species of weeds. Often we have quite detailed data on the crop species, like the interaction coefficients that involve this crop (Guglielmini 2016). An approximation that works in such case is the *focal species approximation* (FSA) (Fort 2020a, 2020b).

Since we are not interested in the yield of each individual weed species, *we can treat all of them as a single species*, and this implies to take the interspecific coefficients between species different from the focal species *k* equal to −1 (remember that *a_ii_*=−1). Thus, one uses the *focal interaction matrix*:

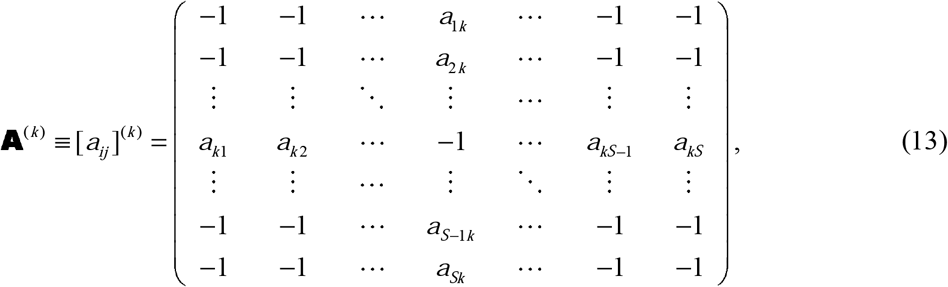

where the *a_ik_* denotes the interaction coefficient of the focal species over species *i* and *a_ik_* the interaction coefficient of species *i* over the focal one. The recipe is then to use matrix (13) to solve Eq. (5) and only take into account the relative yield for the focal species 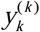 (while neglecting the values 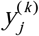 for *j* ≠ *k*). That is, from a matrix of the form (13) we obtain just one relative yield for the focal species 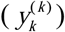.

Figure 2 compares the results of using this approximation with those when using the full set of LLVGE (filled small circles) for four different experiments, three of them involving plants and one for algae.

**Figure 2.**
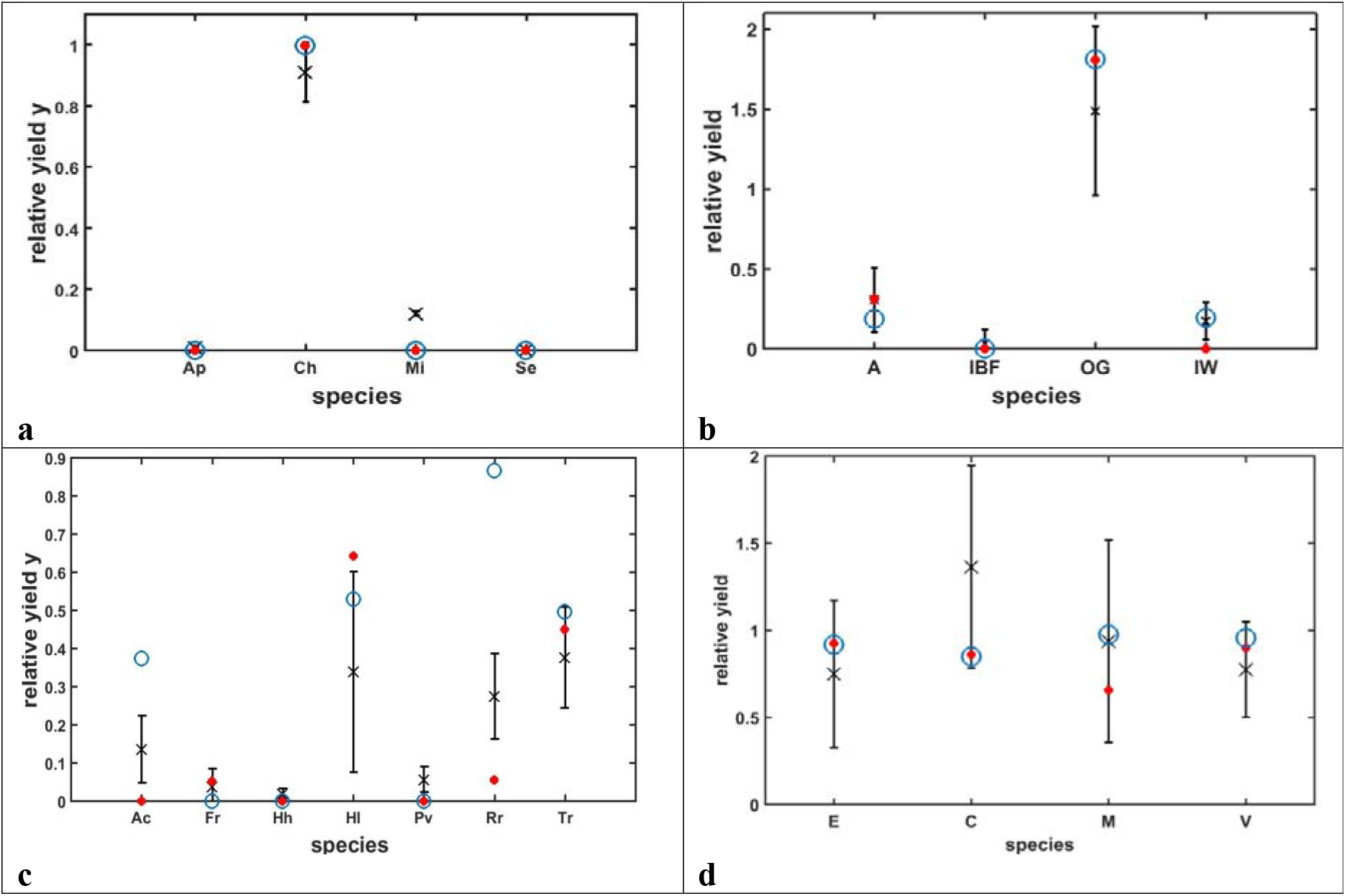
The focal species approximation compared with the full set of LLVGE. Experimental relative yields with their error bars (crosses), predicted values by the LLVGE (red filled circles) and the corresponding values produced by the focal species approximation (open blue circles): a) Algae (Huisman *et al.* 1999). b) Plants A-IBF-OG-IW mixture (Picasso et al. 2008). c) Plants (Roxburgh & Wilson 2000). d) Plants, Area A (Rees *et al.,* 1996).

## 4 CONCLUSION

Therefore we conclude that in cases of incomplete information on the LLVGE parameters:

i. Provided competition interactions among species is dominant, we can predict *aggregate* or *mean* relative competition indices, via de MFA, with an accuracy inversely proportional to the experimental variance of the experimental measurements.
ii. It is possible to predict the yield of a given particular species when we just know the interaction coefficients involving this focal species, by means of the FSA, with accuracy comparable to the one obtained when using the full set of LLVGE.

## Footnotes

1 Thereafter, we will omit the * for all yields in the understanding that they are yields in equilibrium.

## REFERENCES

Andrewartha, H. G., and Birch, L. C. (1954) The distribution and abundance of animals. University of Chicago Press, Chicago.

Beckage, B., and Gross, L. J. 2006. Overyielding and species diversity: what should we expect? New Phytologist, 172, 140–148.

Fort, H. 2020a Ecological Modelling and Ecophysics: Agricultural and environmental applications (IOP ebooks), IOP, Bristol, UK.

Fort, H. 2020b Making quantitative predictions on the yield of a species immersed in a multispecies community: The focal species method. Ecological Modelling 430, 109108.

Fort, H. 2018a On predicting species yields in multispecies communities: Quantifying the accuracy of the linear Lotka-Volterra generalized model. Ecological Modelling 387, 154–162.

Fort, H. 2018b Quantitative predictions from competition theory with an incomplete knowledge of model parameters tested against experiments across diverse taxa. Ecological Modelling 368, 104–110.

Fort, H. & Segura, A. 2018 Competition across diverse taxa: quantitative integration of theory and empirical research using global indices of competition. Oikos 127, 392–402.

Gaston, K. J. (1994) Rarity. Chapman & Hall, London.

Guglielmini, A.C., Verdú, A.M.C., Satorre, E.H. 2017 Competitive ability of five common weed species in competition with soybean. Int. J. of Pest Management 63, 30–36.

Halty, V. et al. (2017) Modelling plant interspecific interactions from experiments of perennial crop mixtures to predict optimal combinations, Ecological Applications 27, 2277–2289.

Huisman, J., Jonker, R.R., Zonneveld, C. & Weissing, F.J. 1999 Competition for light between phytoplankton species: experimental tests of mechanistic theory. Ecology 80, 211–222.

Kirwan, L. et al. (2009) Diversity-interaction modeling: estimating contributions of species identities and interactions to ecosystem function. Ecology 90, 2032–2038.

Krebs, C. J. (1978) Ecology: the experimental analysis of distribution and abundance. 2d ed. Harper & Row, New York.

Lotka, A. J. 1925 Elements of Physical Biology. Williams and Wilkins, Baltimore.

Marcelis, L.F.M., Heuvelink, E. and Goudriaanc, J. (1998) Modelling biomass production and yield of horticultural crops: a review. Scientia Horticulturae 74, 83–111.

Morin, P.J. (2011) Community Ecology. Wiley-Blackwell, Chichester, UK.

NASS-SMB (2012) The Yield Forecasting and Estimating Program of NASS, Statistical Methods Branch (SMB), Statistics Division, National Agricultural Statistics Service, U.S. Department of Agriculture, Washington, D.C., NASS Staff Report No. SMB 12-01.

Pastor, J. 2008 Mathematical ecology of populations and ecosystems. Wiley-Blackwell.

Picasso, V., Brummer, E. C., Liebman, M., Dixon, P. M.& Wilsey, B. J. 2008 Crop species diversity affects productivity and weed suppression in perennial polycultures under two management strategies. Crop. Sci. 48, 331.

Rees, M.P. et al.1996. Quantifying the impact of competition and spatial heterogeneity on the structure and dynamics of a four-species guild of winter annuals. Am.Nat. 147, 1–32.

Roxburgh, S.H. & Wilson, J.B. 2000 Stability and coexistence in a lawn community: mathematical prediction of stability using a community matrix with parameters derived from competition experiments. Oikos 88, 395–408.

Sousa, W. P. 1984. The role of disturbance in natural communities. Ann. Rev. Ecol. Syst. 15:353–391.

Vandermeer, J.H. 1989 The Ecology of Intercropping. Cambridge University Press, Cambridge.

Volterra, V. 1926 Fluctuations in the abundance of a species considered mathematically. Nature118, 558–560.

Wilson, S. D. & Keddy, P. A. 1986 Species competitive ability and position along a natural stress/disturbance gradient. Ecology, 67, 1236–1242.

de Wit, C. T. 1970 Proc. Adv. Study Inst Dyn. Numbers Popul. 269.

